# A database and comprehensive analysis of the algae genomes

**DOI:** 10.1101/2021.10.30.466624

**Authors:** Chengcheng Shi, Xiaochuan Liu, Kai Han, Ling Peng, Liangwei Li, Qijin Ge, Guangyi Fan

**Affiliations:** BGI-Qingdao, BGI-Shenzhen, Qingdao, 266555, China

## Abstract

Algae characterize their high diversity, taxonomy and morphology for wide-used studying the plant origins and terrestrialization, as well as multicellular evolution. Due to the genome assembly challenge of algae caused by symbionts with microbiome, the published algae genomes are relatively less than the terrestrial plants. Here we comprehensively collected and re-annotated 191 available algae genomes distributed in nine major lineages. We systemically investigated the genome features including genome size, assembly continuity and integrity, GC content, abundance of repetitive sequences and protein-coding gene number. We construct the phylogenetic trees using 193 algae genomes, which is consistent with the well-known evolution path that Glaucophyte is the most ancient, going through eight lineages, and finally evolved to terrestrial plants. We also examined the Horizontal Gene Transfer (HGT) genes distribution in algae genomes and provides a substantial genomic resource for functional gene origins and plant evolution.

## Introduction

Algae are a large group of photosynthetic eukaryotic organisms distinguished from land plants, being of vast diversity in terms of taxonomy, morphology and genetic features. Most of them are aquatic organisms, and small fraction of them inhabit in soil, desert, rocks, vegetation, even fur of animal. They exhibit adaptability to various environmental conditions including halotolerance, thermotolerance, freezing tolerance and acid tolerance, consequently distributing worldwide. The red algae *Galdieria sulphuraria* can grow at pH 0 to 4 and temperatures up to 56°C, close to the upper temperature limit for eukaryotic life [1]. *Emiliania huxleyi* as the first haptophyte reference genome, high genome variability was found, which enforced its capacity to thrive in different habitats ranging from the equator to the subarctic, and to form algal blooms under a wide variety of environmental conditions[2]. In addition, green algae *Dunaliella salina* [3], *Picochlorum SENEW3* [4] and *Picochlorum renovo* [5] is highly tolerant to salinity; Green algae, *Trebouxiophyceae sp. KSI-1* is highly tolerant to oxidative stress [6]; and *Coccomyxa subellipsoidea* tolerate to extreme temperature isolated from Antarctica, which provide us appealing models to investigate metabolic processes involved in stress responses in algae.

Algae is polyphyletic and includes multiple distinct clades originated form endosymbiosis. Among of them, green algae are profoundly important organisms composed of Chlorophyta and Charophyta, including more than 10,000 species, inhabiting almost every environment. Red algae (Rhodophyta) is one of the oldest and the largest categories of algae, which is more than 7,200 species [7-10]. Many red algae synthesize unique polysaccharides, which contribute to their ecological success and are of significant biotechnological and industrial application[11]. Heterokont (stramenopiles), as a major line of eukaryotes, includes around 10,997 species (summarized from NCBI Taxonomy database https://www.ncbi.nlm.nih.gov/taxonomy). Algae species in Heterokonts are mainly colored groups represented by diatoms, brown algae, golden algae, and yellow-green algae, covering different smaller groups which disperse in diverse clades. Haptophytes may be famous for destructive toxic algal blooms[12] and their contribution to global carbon fixation[13]. In addition to the more familiar algae mentioned above, there are also some studies focus on those algae that we don’t usually pay attention to. For cryptophyceae, euglenida and chlorarachniophyta, endosymbiosis is an interesting study topic and they all acquire plastids from eukaryotic algae [14-16]. The complicated genetic source of course leads to genetically complex genome composition: mitochondrial, plastid, “master” nuclear, and residual nuclear genome of secondary endosymbiotic origin, so-called “nucleomorph” genome [17]. Meanwhile, cryptophyceae genomes have been independently reduced and compacted, such as *Hemiselmis andersenii*, which completely loses intron in nuclear genome and has less gene number [18], the similar genome reduction has been found in *Cryptomonas paramecium* [19] with gene loss, but spliceosomal introns are present in the nucleomorph genome of *Chroomonas mesostigmatica* and it has been reported that the complete loss of spliceosomal introns occurred within the *Hemiselmis* clade [17].

Genomics research provides us with a specific perspective of algae origin and evolution. Whereas, the available algae genomes are just a tip of the iceberg in algae species. Compared to the large number of species that this group contains, the amount of data available today is far from enough to make clearly explanations. There are several factors hamper the assembly and research of algae genomes. Algae are commonly symbiosis with various bacteria, fungal, lichens, and corals, which make it difficult to exclude contaminations in experiment. In addition, the huge genome is another characterize of dinoflagellate. The relatively lagging progress of valuable research seem to be inseparable from technical difficulties. Enormous genome size has brought ineluctable challenges to handling massive data and acquiring high-quality assemblies at present. A few years ago, the assemblies of some gymnosperms with large genomes more than 20Gb were obtained through high-throughput sequencing, but most of them were relatively fragmented as the scaffold N50 only with tens of kb [20-22], even though few can reach over 200kb using data from multiple platforms [23]. Imaging a dinoflagellate with a genome size of more than 30Gb, for instance *Alexandrium*, the sequenced data required to assemble the whole genome probably exceed 2Tb (∼80X). Moreover, considering the assembly completeness, such large genome is generally recommended to be sequenced with third generation sequencing technology. The large order of magnitudes together with the potential complex sequence characteristic call for extremely high requirements on software algorithms and computing resources. In this work, we collected all the available alga genome data, and re-annotated the protein-coding genes and repetitive sequences. We systematically compared the genome characters, and investigated the highlighted interests of the main algae phyla.

## Results

### The landscape of sequenced algae genomes

To date, a total of 191 algae genomes are publicly available, covering most of the known algae phyla including green algae (111 Chlorophyta and 6 Charophyta), 34 Heterokonts (e.g. diatom), 12 Rhodophyte, 10 Dinoflagellates, 6 Haptophyte, 5 Cryptophyceae, 1 Glaucophyte and so on (**Fig .1a**). Among the sequenced algae genomes, 57.59% are green algae, followed by the Heterokonts (17.80%), red algae (6.28%), Dinoflagellates (5.23%) and Haptophyte (3.14%). All of the five mentioned groups make up of 91.10% (174) of the published algae genomes (191) (**Fig. 1a**). As the group with the largest population in algae, green algae are the most well studied and has the largest number of sequenced genomes to date. Of the 117 species green algae been sequenced and their genomes been assembled, vastly diverse of life forms were observed, ranging from unicells to large and complex multicellular or siphonal life forms (**Fig. 1b**). The unicellular green alga *Ostreococcus tauri* (Prasinophyceae) is the known smallest free-living eukaryote with a genome size of 12.56 Mb [24], while on the other hand, the siphonous macroalgal *Caulerpa lentillifera* could reach meters in size, and probably is the largest single cell on earth with a genome size of ∼29 Mb [25]. Although Chlorophyta dominated the green algae, most of the sequenced genomes are from three classes (Chlorophyceae, Trebouxiophyceae and Mamiellophyceae, accounting for 92.8%), leaving large fractions of species from the other classes to be studied**(Fig. 1b)**.

**Figure 1.**
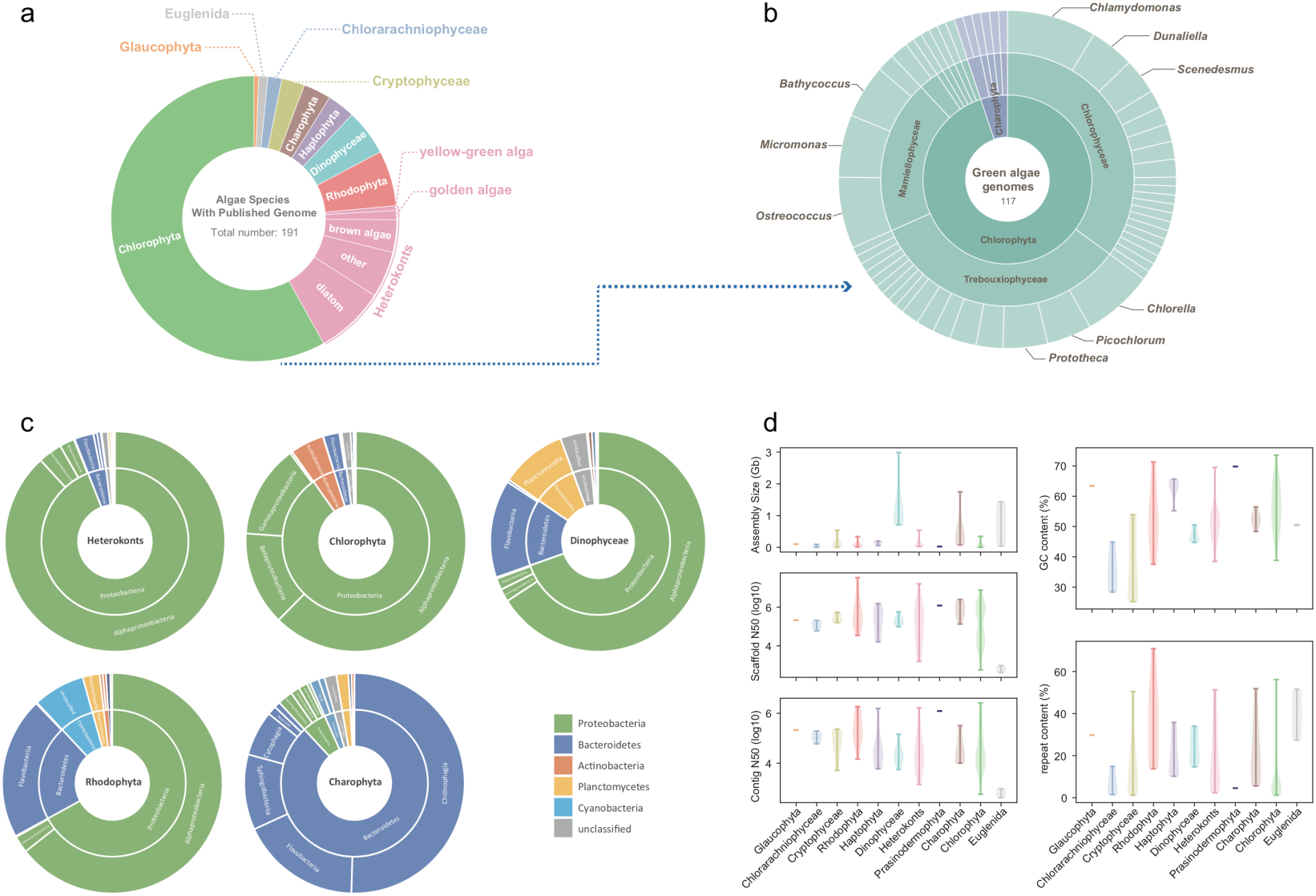
Statistics of sequenced algae genomes. a) The number and phylogeny of public algae genome. The circle was classified by phylum. b) The number and phylogeny of public green algae. From the inner circle to the outer circle were classification of phylum, class and genera. Names of genera with species number greater than three were indicated. c) Contaminants categories distribution in each algae phylum. d) The assembly size, scaffold N50, contig N50, GC content and repeat content of algae genomes. Colors indicate different algae.

Prokaryotic genome contamination in algal genomes could harm the study of algae, for example it can greatly increase the false positive rate when detecting horizontal gene transfer events. To assure the results of this study, all potential prokaryotic genome contaminant sequences were filtered out before downstream analysis. A total of 10,020 sequences from 84 algal assemblies were identified as contamination and discarded, with a total length of 83.7Mbp. The potential contamination rare could be as high as 25.61%, as observed in *Halamphora sp. AAB* in the current study **(Table S1)**. In addition, different bacteria phyla contribute to the major fraction of contamination of different algae assemblies. For example, we found that Proteobacteria accounted for most of the contamination in Heterokonts and Chlorophyta, while in Dinophyceae and Rhodophyta, besides Proteobacteria, a small fraction of Bacteroidetes contamination is also observed, and The main contaminant bacterial phylum in Charophyta is Bacteroidetes (mostly from genome of *M. endlicherianum*) **(Fig. 1c)**. However, it is important to note that due to differential experimental processes, contamination observed in genome sequencing data might not be used to infer a symbiotic relationship of the observed bacteria and algae.

Features, including genome size and GC content, and qualities of the assembled algae genomes were assessed., The genome size of most the algae sequenced (e.g. Chlorophyta and Rhodophyta) were arount ∼100 Mb, except for Charophyta, Dinoflagellate and Euglenida, the genome size of which were greater than 400Mb (**Fig. 1d**). An uneven distribution of the completeness of the algae assemblies was observed, with Rhodophyta assemblies showing the highest average scaffold N50 of 1 Mb and contig N50 of 400 Kb (**Fig. 1d**), suggesting high quality of these genomes. Differential GC ratio of these genome assemblies were observed with Glaucophyta and Haptophyta having the highest average GC ratio of greater than 60% (**Fig. 1d**).

### Phylogeny and Evolution of algae

Due to their diverse and distinct morphology and taxonomy, the evolutionary trajectory of different algal groups have been elusive for long. Lots of scientific efforts and attempts have been made to classify different algae groups and reconstruct their phylogenetic relationships based on either ultrastructural features of cell and plastids, or their genome sequence features, or a combination of both. According to Carrington and colleagues, “the story of algae evolution is tied to how they acquired photosynthetic organelles or plastids”(ALGAL EVOLUTION n.d.). It is now acknowledged that the single event of ancient primary endosymbiosis led to the divergence of three algae lineages, including Glaucophyta, Rhodophyta, and Chlorophyta, while several other independent events of complex endosymbiosis by different host eukaryote formed other algae lineages, such as Chlorarachniophyta, Euglenophyta, Heterokontophyta, and Haptophyta. Although there is a consensus regarding what determines algae origin, detailed phylogenetical track of different lineage is still challenging and remains to be discussed.

With all available genome sequences of 193 algae species as well as two genomes from land plant (*Arabidopsis thaliana* and *Oryza sativa*), we reconstructed the phylogenetic relationship based on 255 conserved genes (from BUSCO eukaryota_odb10 dataset) (**Fig. 2a and 2b**). The evolutionary track of each busco gene was inferred using IQ-TREE software with best suited substitution model after filtering out gappy sites and fragmentary sequences, as well as abnormally long branches, and leaves with low bootstrap support were deleted and collapsed. Species trees were estimated using ASTRAL software by summarizing gene trees (**Fig. 2c and 2d**). Two distinct algae categories were observed based on their phylogeny, one of which was grouped with Rhodophyta and the other belonged to the lineage developed from Chlorophyta. The Dinophyceae, commonly thought to have a complex evolutionary line including secondary endosymbiosis events with green or red algae, and tertiary endosymbiosis with haptophyta, was clustered into the rhodophyta clade and jumbled with one chlorophyta species as well as one heterokonts species. The algae group Chlorarachniophyceae was indicated as a sister clade of Heterokonts and unexpectedly clustered into the clade of Rhodophyta, which was not in accordance with the perception of a green algae derived origin. Our results showed a clear separation of the green algae group and the Charophyta, and also demonstrated potential close relationships of land plants to charophyta species. What’s more, species *Prasinoderma colonial* (a recently novel phylum) also formed a single branch before the split of Chlorophyta and Streptophyta.

**Figure 2.**
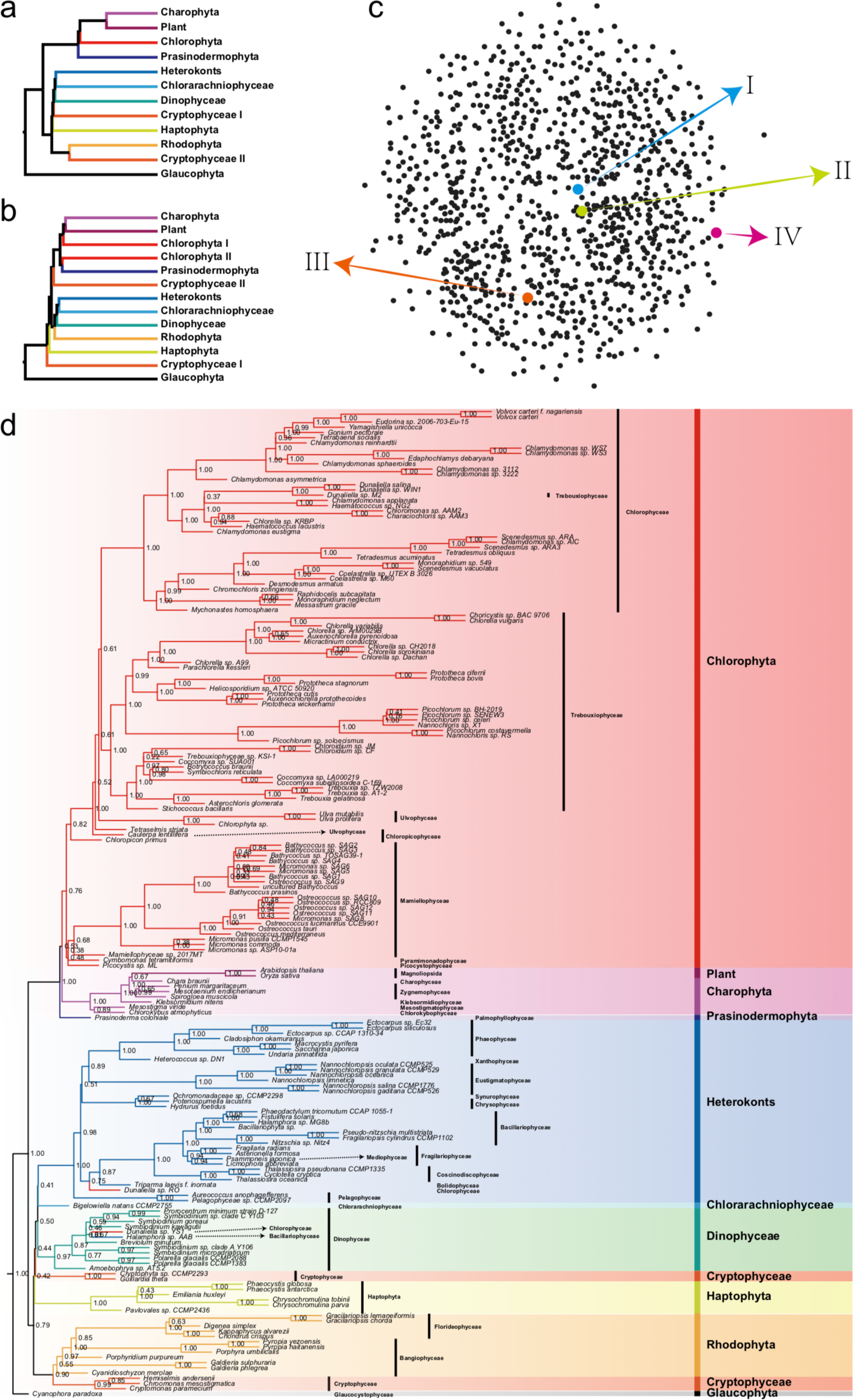
Phylogenetic relationships of algae based on summary method of conserved BSUCO sequences. a) Main phylogenetic topology of algae phylum inferred by busco proteins and combination of proteins and nucleotide sequences, which was well supported by random sampling. b) phylogenetic topology of algae phylum inferred using busco nucleotide sequences. c) Multidimensional scaling based on un-weighted Robinson-Foulds distances, black dots represent species trees summarized from random sampling gene trees, I busco combination tree, II random sampling consensus, III busco peptide tree, IV busco nucleotide tree. d) Species tree summarized from combination of busco protein and nucleotide, branches leading to different algae groups were very short.

### Diversity of transposable elements in algae

Transposable elements (TEs) is a type of important genomics repetitive elements that can move to new sites of the genome and thus might disrupt normal gene functions and alter genome architecture. In algae, relatively lower TEs ratios ranging from 5%∼40% were observed than terrestrial plants, which generally contain around 50% of repetitive sequences (**Fig. 1d**). A closer look at of the TE types indicated that the type of LINE and LTR in class I and DNA in class II made up the dominant proportion, and tandem repeat was also distinct in all species. The proportion of the class I repeat sequence in these genomes is more constrained than class II (**Fig. 3a**). What’s more, the green algae was observed to have more diverse tandem repeat contents than other algae, and the genome size and repeat sequence ratio of dinoflagellates are huge in size remarkably. Within species analysis revealed that LINE in seven green algae while LTR in the red algae genomes were the prominent types of TE, respectively (**Fig. 3b**). Phylogenetic trees were constructed using RT region of TEs. In contrast to Chlorophyta, of which their genomes were not rich in TEs, a large number of LINE and Gypsy sequences were found in Charophyta, while Gypsy and Copia are relatively abundant with almost on LINE in Rhodophyta. It is noteworthy that LINE in Charophyta seems to have independent origins with other green plants.

**Figure 3.**
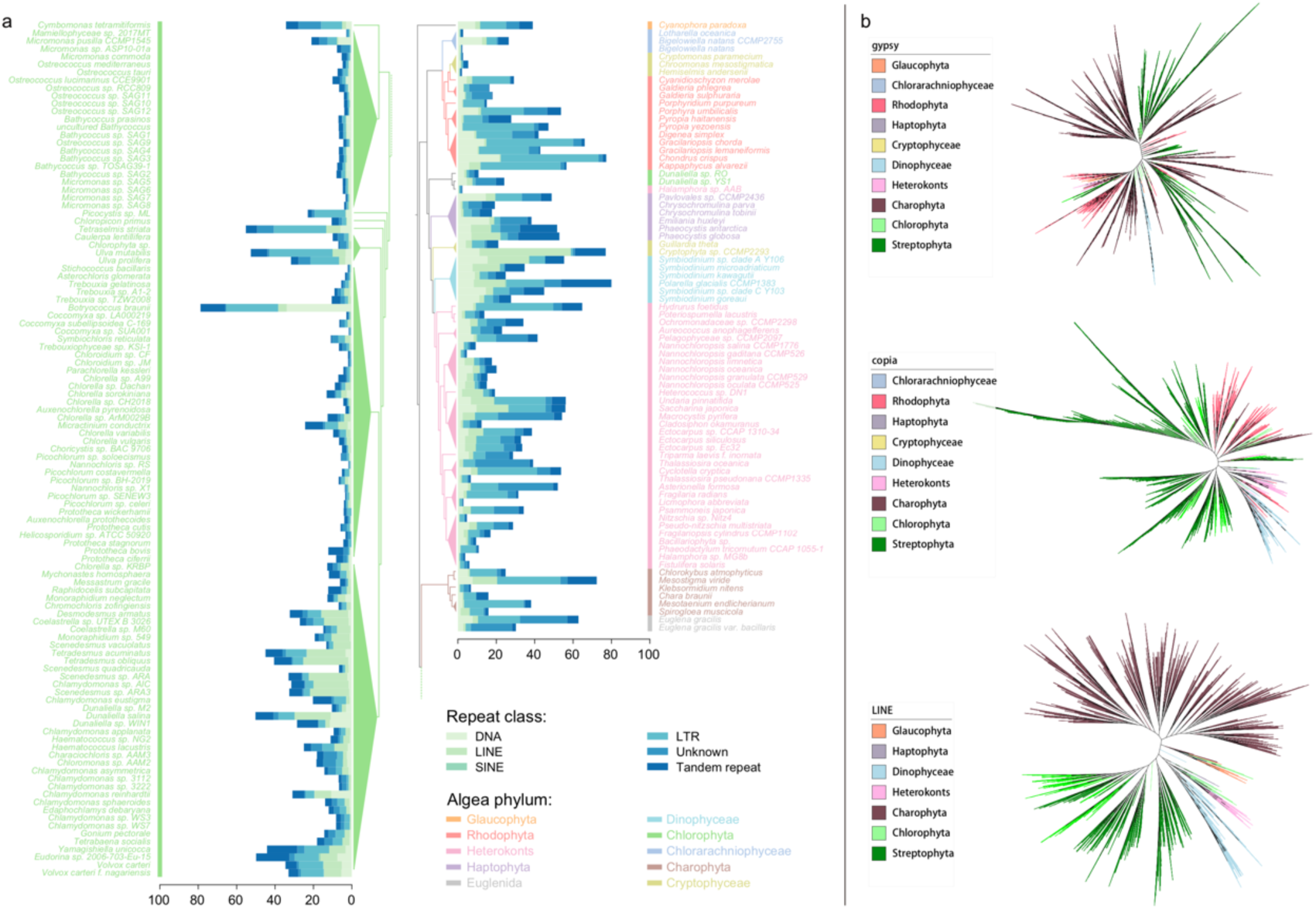
Summary and phylogeny of TE types in algae genomes. a) The proportional distribution of TE types (DNA, LINE, SINE, LTR and others) and tandem repeat in algae genomes. The lines on the left and right indicates the remarkable LINE and LTR content in some green algae and red algae genome, respectively. b) The TE phylogeny of LINEs and LTRs in green alage and red algae respectively.

### Horizontal gene transfer

As horizontal gene transfer (HGT) is one of the important factors for the evolution of bacteria [26], HGT also helps algae to acquire new genes from the environment to improve their metabolic diversity and environmental adaptability [27, 28]. HGT in green algae and red algae are well documented, with some of them being related to carbohydrate metabolism, osmolyte regulation, sulfate scavenging and cell cycle control [27], some of them about energy shuttle and arsenic detoxification[28], and some facilitating algae to adapt to extreme environments [1]. In order to compare HGT genes between different algae, 172 algae,including 7 species of Charophyta, 1 species of Chlorarachniophyceae, 99 species of Chlorophyta, 2 species of Cryptophyceae, 8 species of Dinophyceae, 1 species of Euglenida, 1 species of Glaucophyta, 6 species of Haptophyta, 34 species of Heterokonts, 1 species of Prasinodermophyta and 12 species of Rhodophyta, were selected and their genomes carefully evaluated.

Except for a few algae species, the fractions of HGT genes in most algae are relatively low (**Fig. 4a**). Functional annotation of the HGT genes against KEGG database revealed a widespread KEGG pathway distribution of algae from Dinophyceae, which might be caused by the enormous genes and huge genome size of Dinophyceae. Similarly, the HGT genes of *Halamphora sp. AAB* and *Hydrurus foetidus* of Heterokonts and *Digenea simplex* of Rhodophyta also showed wide distribution. That was consistent with the relatively high HGT gene percentage (**Fig. 4a**) and indicated that these algae might have more active genetic exchange with environments. Closer look at of the pathways revealed that Carbon metabolism, biosynthesis of amino acids, metabolic pathways and biosynthesis of secondary metabolites showed highly consistent patterns among certain algae species. Form the category of algae, although species of Chlorophyta are more than half, there is no one with enormous HGT genes. And the same result cloud observed from the percentage of HGT genes, that Chlorophyta is at a quite low level (**Fig. 4a**). Consistent with previously published documents, we also found that HGT genes of *Porphyridium purpureum* were associated with photosynthesis[29] and HGT genes of *Pyropia yezoensis* were associated with glycine-serine- and-threonine metabolism, metabolic pathways, peroxisome and nitrogen metabolism [30].

**Figure 4.**
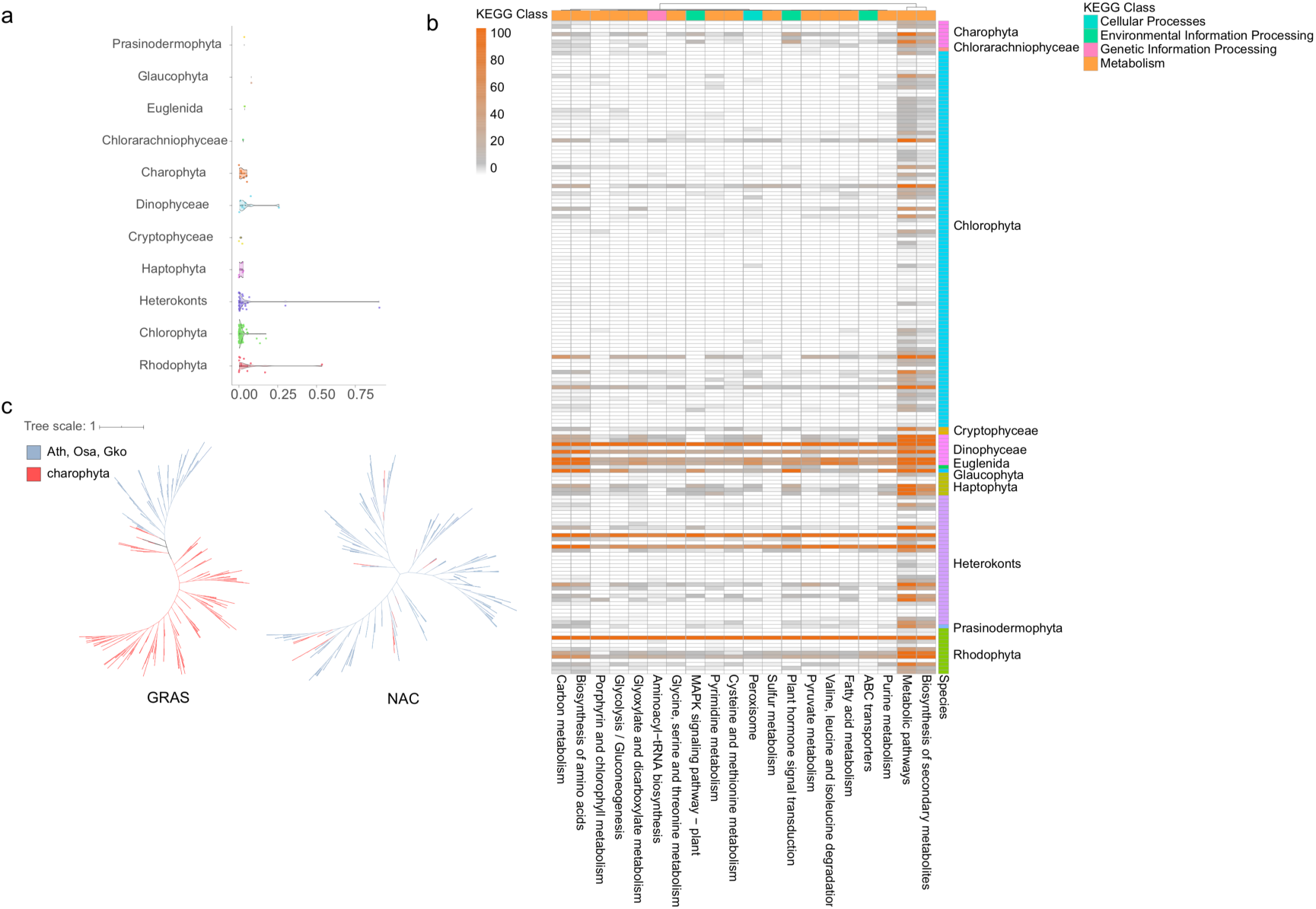
Classification of HGT genes and phylogeny of TF. a) The HGT gene percentage of algae. b) The heatmap of KEGG pathways of 172 species of algae. c) The phylogeny of GRAS and NAC genes.

### The origin of land plants and terrestrialization

The terrestrialization of green plants is a pivotal to understand the evolutionary trajectory of land plant. The closest relatives of land plants (embryophytes) are charophytic algae with both of them constitute Streptophyta. As a morphologically diverse group encompassing unicellular and structurally complex multicellular, Charophyte algae includes six distinct major lineages: Mesostigmatophyceae, Chlorokybophyceae, Klebsormidiophyceae, Zygnematophyceae, Charophyceae, and Coleochaetophyceae [31]. Genome study of Charophyte could provide insight into the origins of land plants and the underlying molecular mechanism of plant terrestrialization. It has been reported that transcription factor (TF) genes that could improve the resistance to biotic and abiotic stresses in land plants originated or extended from the common ancestor of Zygnematophyceae and embryonic plants. A particularly large number of expanded gene families including GRAS, HDKNOX2, BBR/BPC, NAC, LOB, bZIP, basic-helix-loop-helix (bHLH), WRKY, and ERF families [32, 33] have been observed and studied. The bursts of gene family expansion were evident coincides with the emergence of various charophyte lineages [33]. It was proved that genes (i.e., GRAS and PYR/PYL/RCAR) that increase resistance to biotic and abiotic stresses were gained by horizontal gene transfer (HGT) from soil bacteria [32]. Phylogenetic trees were constructed using TF genes across 16 representative chlorophyta, charophyta, *Arabidopsis thaliana, Oryza sativa*, and *Ginkgo biloba*. We can find that GRAS and NAC genes only in charophyta and land plants, coinciding with previous report that the common ancestor of the charophyta and land plants has acquired GRAS and NAC genes after split from chlorophyta **(Fig. 4c)**. GRAS can be clustered to a whole clade, whereas NAC genes could apparently cluster to three clades, suggesting the differential evolution patterns of these two TF gene families.

## Discussion

Without relatively complete genome information, it’s hard to reveal a clear picture that how algae evolved and how they became what they are now. Another problem that obscures genome assembly is sequence contamination by bacteria, because commonly algae is in symbiosis with bacteria. Of course, some studies have separated algae sequences from contamination by mapping sequence to NR library [11, 30, 34-36]. But if we could separate them experimentally before sequencing or develop some powerful and robust algorithm to distinguish them, maybe the assemblies will be more accurate and the false-positive result of downstream analysis, such as HGT candidate genes identification, will be decreased dramatically. In fact, there are many software to solve this problem based on the differences in sequence characteristics (GC content, k-mer frequency, genome abundance, tetranucleotide frequency etc.) of eukaryotic and prokaryotic organisms, such as EukRep[37], Kraken2[38] and k-means[39] algorithm for clustering. In addition, trio-binning strategy which is used for haploid assembly, is also a feasible way to remove variable exogenous contamination through the batch effect of sequencing data, and obtain a relatively pure algae genome. HAST[40], a recently published software, may realize this vision if there is suitable data. Alternatively, we may open mind more widely, change our mind and turn the question of ‘how to separate algae genome with contamination’ to ‘what species do these sequences belong to, no matter it is algae or bacteria’, that means we should change from common genomic mind to meta genomic mind. And when we think it as meta genomes, binning software, such as MetaWrap[41], cloud use multiple algorithms to sperate the sequences into different bins. And with the assistance of 16S/18S identification, we could not only get the species information of the algae and the many bacteria that exist in the same habitat, but also could help us to understand collaboration and cooperation genetically and/or ecologically between algae and microbes, even could supply extra information to separate algae squences from microbes’. That will facilitate the studies of algae symbiotic organisms and show a whole picture of the genetic exchange between algae and symbiotic bacteria.

It is worth mentioning that transcriptome data is one important part in genomics research, for the transcriptional regulation of related genes can be observed. Comparative genomics and transcriptomics were used to explore the evidence that remains in genomes during adaptive evolution as well as to excavate its mechanisms. Comparative transcriptomics analysis of *Picochlorum renovo* under different salinity experimental conditions confirmed previously reported genes governing proline metabolism which is involved in the high-salt response, and also observed a series of previously unreported halo-responsive genes including ppsA, ppsC, pks1, pks15, iput1, cerk, rad54 and dmc1 [5]. Intriguingly, there is a striking overlap between the thermotolerance and halotolerance transcriptional rewiring in Picochlorum SE3 [42]. Combining transcriptional regulation network analysis of microRNA system, authors comprehensively illustrated a biochemical complementarity between the *F. kawagutii* and coral genome, and constructed a mechanisms schema of symbiosis and cargo transport [43]. Genomic data is the foundation, and the combination of multiple omics data will be necessary for studying biological functions, especially for dinoflagellate with complex habitats. Recently, single cell transcriptome technology was widely used in studies to explore the dynamic development of cells. In the study of soft coral Xenia endosymbiotic cell lineage, through single-cell sequencing of symbiotic and non-symbiotic cells distinguished from different cell stages, the researchers constructed an endosymbiotic system of corals and algae [44]. This kind of project designment is very instructive, especially in the study of dinoflagellates symbiotic. Some algae are directly related to the harmful ecological phenomenon, algal blooms. From the perspective of ecological research, the combined application of multiple omics and new technologies is an inevitable trend. Furthermore, multidimensional analysis with integration of genomics, transcriptomes, proteomics and utilization of the cutting-edge single-cell technology, as well as evidence from comparison of multiple species or even different groups, would be helpful for building a robust model.

## Method

### Evaluation of algae contamination

The specific method was as follows, genomes were compared to NCBI Nucleotide database using BLAST; the number of hints refer to prokaryotic or eukaryotic at each site were counted. If the number of prokaryotic hints at a certain site were greater than the number of eukaryotic hints, this site was considered as a contaminated site; if the total number of contamination sites in a sequence is greater than 50% of the total length of the sequence, this sequence is considered to be a contaminated sequence.

### Identification of horizontal gene transfer

We firstly aligned the algae genes to NR database [45] using Blastp [46] (v2.6.0, with parameters ‘e-value 1e-5’). Then for each gene, the top 50 hits with the smallest e-values were retained. After being assigned to taxonomy, each gene cluster was checked independently using the follow criteria: if over 90% of hits, including the best hit, were assigned to different superkingdom taxonomies from that algae, this gene was defined as a candidate HGT gene.

### Algae evolution and terrestrialization of green plants

The evolutionary track of each BUSCO gene was inferred using IQ-TREE [47] software with best suited substitution model after filtering out N sites, fragmented sequences. The abnormally long branches as well as branches with low bootstrap (<50) support were further deleted and collapsed. Species trees were estimated using ASTRAL [48] software by summarizing gene trees.

